## Supplemental Figures for "Neural circuits underlying habituation of visually evoked escape behaviors in larval zebrafish"

### Supplementary materials

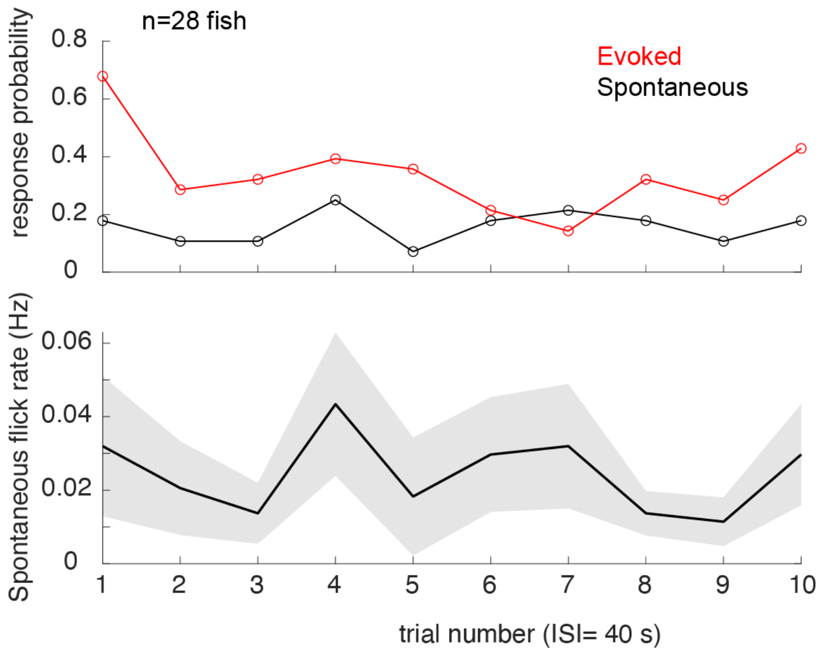


**Supplementary Figure 1. Spontaneous tail flicks occurred at a low rate, which did not change throughout the experiment.** Top panel shows average response probability of stimulus-evoked (red, calculated for each trial across 28 fish) and spontaneous tail flicks calculated in a same-length window prior to stimulus presentation (black). For each fish a response was registered as 1 each time there was a tail flick and zero when there was none. The probability of evoked responses declines over trials, whereas it stays stable for spontaneous. Bottom panel shows the average rate of spontaneous tail flicks was similarly stable across trials.

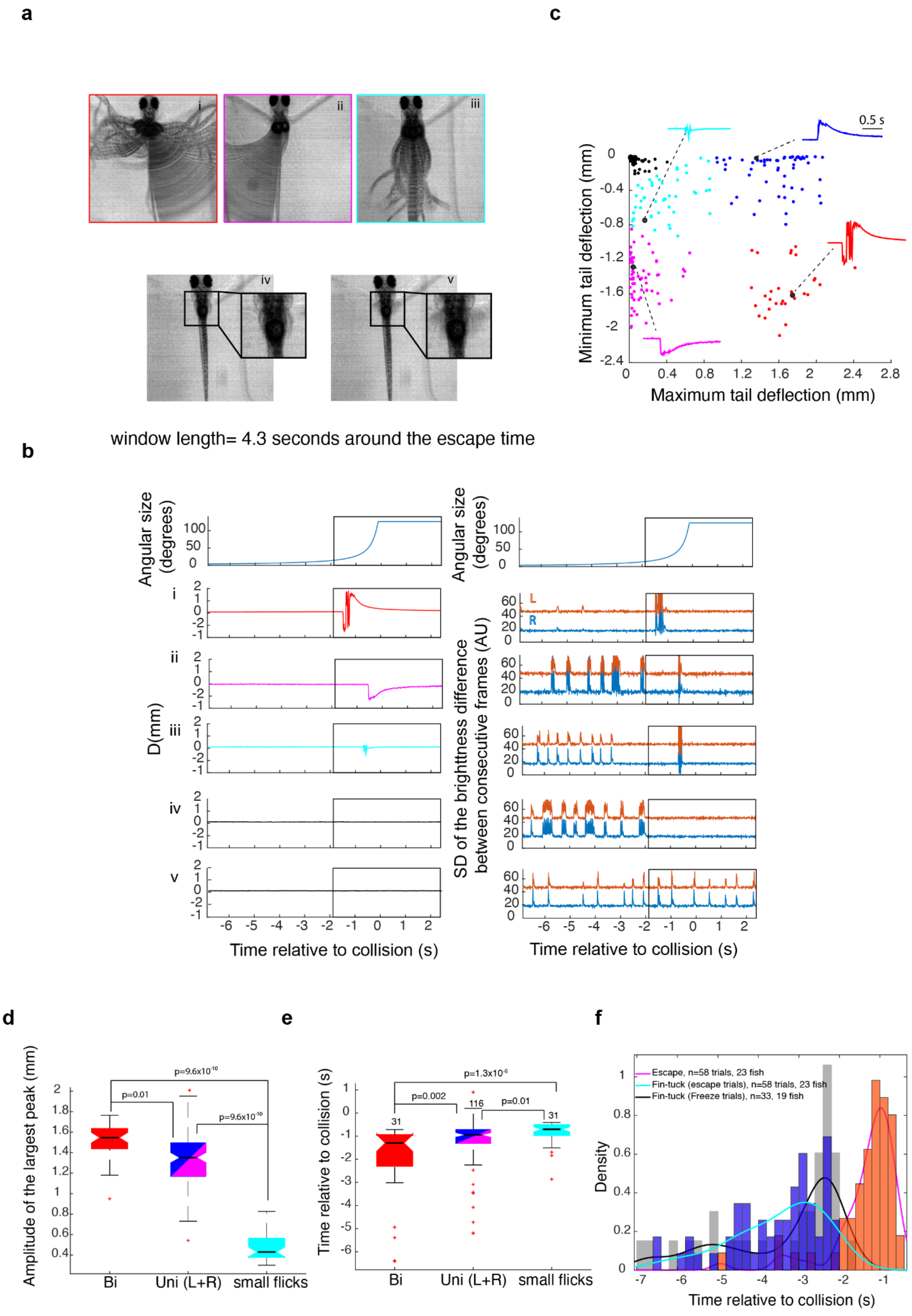
a
i
ii
iii
iv
v
c
0.5 s
Minimum tail deflection (mm)
Maximum tail deflection (mm)
0
-0.4
-0.8
-1.2
-1.6
-2
-2.4
0
0.4
0.8
1.2
1.6
2
2.4
2.8
window length= 4.3 seconds around the escape time
b
Angular size (degrees)
100
50
0
D(mm)
2
1
0
-2
-1
L
R
60
40
20
0
SD of the brightness difference between consecutive frames (AU)
-6
-5
-4
-3
-2
-1
0
1
2
Time relative to collision (s)
d
Amplitude of the largest peak (mm)
p=9.6x10-10
p=0.01
p=9.6x10-10
2
1.8
1.6
1.4
1.2
1
0.8
0.6
0.4
Bi
Uni (L+R)
small flicks
e
Time relative to collision (s)
p=1.3x10-6
p=0.002
p=0.01
31
116
31
1
0
-1
-2
-3
-4
-5
-6
Bi
Uni (L+R)
small flicks
f
Density
Escape, n=58 trials, 23 fish
Fin-tuck (escape trials), n=58 trials, 23 fish
Fin-tuck (Freeze trials), n=33, 19 fish
1
0.8
0.6
0.4
0.2
0
-7
-6
-5
-4
-3
-2
-1
Time relative to collision (s)

**Supplementary Figure 2. Larval zebrafish generate various types of escapes and freezing responses to looming stimuli. a .**Composite video frames corresponding to various response types calculated over a 4.3 s- long window around the time of escape. Insets in panels iv-v focus on a window around the pectoral fins. **b**. Tail and pectoral fin traces corresponding to the examples shown in **a**. The top panels show the time course of the stimulus angular size. The left panels below show the average of tail deflection from the midline and the right panels show the corresponding fin movement traces. Note that escape movements create an artifact in the fin trace (large deflections within the rectangular window. The rectangular window corresponds to the time window where the composite images in panel **a** were calculated. **c**. K-means clustering of response types: response time, amplitude of the positive peak, amplitude of the negative peak and their ratio were used to classify behavior into four different escape types (colored dots) and no escapes (black dots). The panel shows the result of K-means clustering plotted against two of the variables used for clustering data from 420 trials in 42 fish. The stimulus was either presented to the left or right eye, resulting in two clusters of unilateral escapes showing opposite tail flick directions. The insets show the points corresponding to traces shown in panel **b**. Red: large bi-directional tail flicks, magenta and blue: uni-directional tail flicks to one side or the other, cyan: small flicks, black : no escape **d,e**. Amplitude and timing of escape relative to expected collision for different response types, respectively. Data for left and right unidirectional tail flicks were pooled. **f.** The distribution of the timing of escape and fin tucking for escape and freeze trials. Although fin-tucks occurred significantly earlier than escapes, the timing of fin tucks were not significantly different between escape and freeze trials.

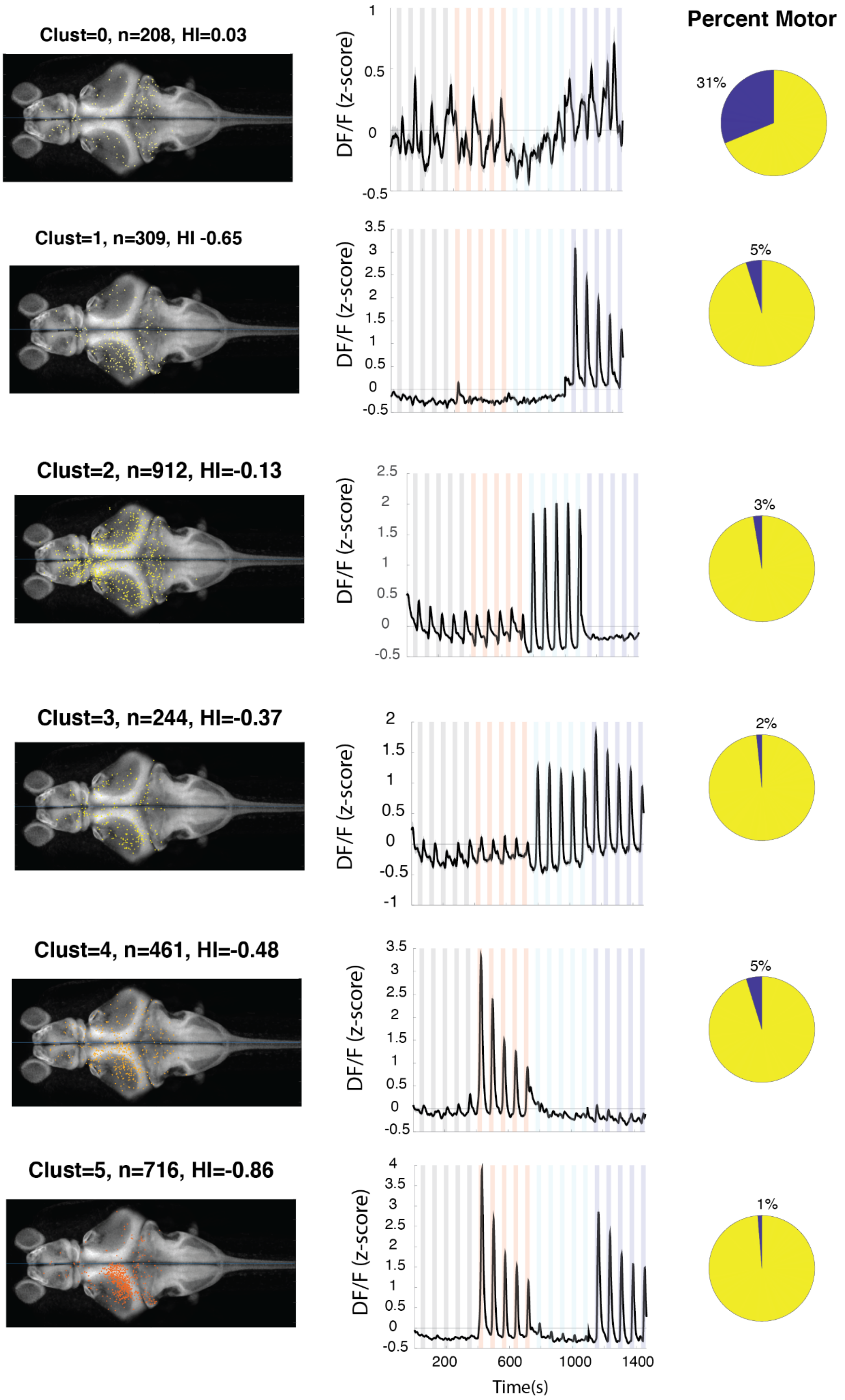

Percent Motor
Clust=0, n=208, HI=0.03
31%
Clust=1, n=309, HI -0.65
5%
Clust=2, n=912, HI=-0.13
3%
Clust=3, n=244, HI=-0.37
2%
Clust=4, n=461, HI=-0.48
5%
Clust=5, n=716, HI=-0.86
1%
DF/F (z-score)
Time(s)
200
600
1000
1400

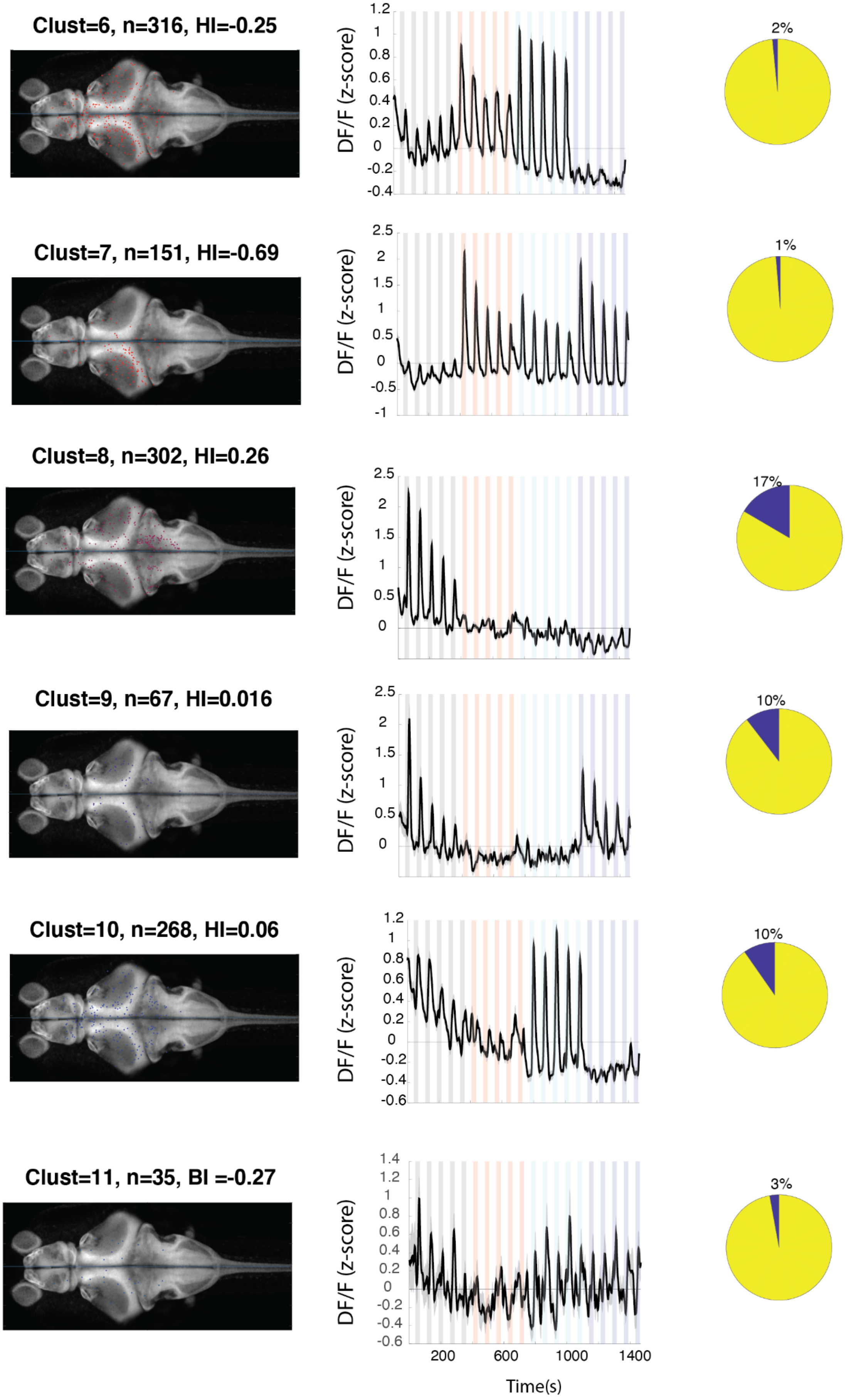
Clust=6, n=316, HI=-0.25
Clust=7, n=151, HI=-0.69
Clust=8, n=302, HI=0.26
Clust=9, n=67, HI=0.016
Clust=10, n=268, HI=0.06
Clust=11, n=35, BI =-0.27
DF/F (z-score)
2%
1%
17%
10%
10%
3%
200
600
1000
1400
Time(s)

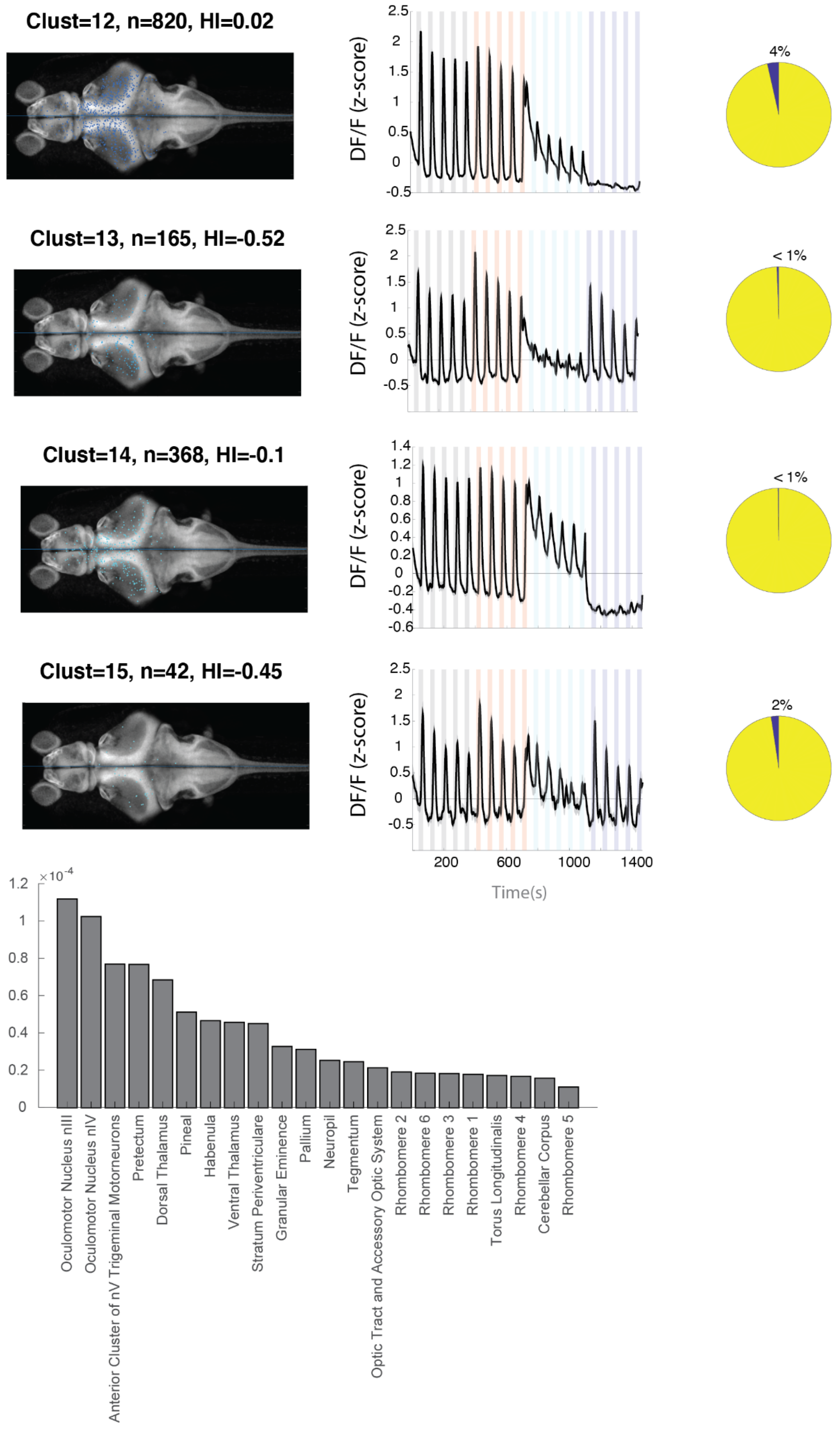

Clust=12, n=820, HI=0.02
Clust=13, n=165, HI=-0.52
Clust=14, n=368, HI=-0.1
Clust=15, n=42, HI=-0.45
DF/F (z-score)
Time(s)
4%
< 1%
< 1%
2%
×10^-4
Oculomotor Nucleus nIII
Oculomotor Nucleus nIV
Anterior Cluster of nV Trigeminal Motorneurons
Pretectum
Dorsal Thalamus
Pineal
Habenula
Ventral Thalamus
Stratum Periventriculare
Granular Eminence
Pallium
Neuropil
Tegmentum
Optic Tract and Accessory Optic System
Rhombomere 2
Rhombomere 6
Rhombomere 3
Rhombomere 1
Torus Longitudinalis
Rhombomere 4
Cerebellar Corpus
Rhombomere 5

**Supplementary Figure 3.** Left columns : distribution of neurons within each cluster across brain regions mapped onto the standard Z-brain**.** Clust: Cluster number, HI: Hemispheric index, n: number of neurons. Middle columns: average response of neurons in all 16 clusters. Right column : percentage of neurons in the cluster that show correlation of more than 0.5 with the motor output. Histogram shows the neuron density across brain regions that were imaged. Regions are shown only if they contained at least 10 cells and were represented by at least 3 fish.
